# Building the Marine Probiotic Platform: Developing a High-Throughput Process to Discover Coral Probiotics for Stony Coral Tissue Loss Disease

**DOI:** 10.64898/2026.01.11.698879

**Authors:** Margaret K. Walter, Blake Ushijima

## Abstract

Stony coral tissue loss disease (SCTLD) is a deadly, waterborne coral disease. Since 2014, SCTLD has spread throughout Florida’s Coral Reef and is now confirmed in 28 Caribbean countries and territories. The causative agent(s) remain unknown; however, pathogenic bacteria are implicated in disease progression. In contrast, a beneficial microbe (a probiotic) with antibacterial activity isolated from a disease-resistant coral has arrested SCTLD transmission and progression during laboratory and field trials. Due to host and environmental specificity, more probiotics sourced from vulnerable coral species and different SCTLD-impacted regions are needed. Conventional methods used for probiotic discovery include plating samples for single colonies, streaking for purification, and then testing strains for antibacterial activity with drop culture assays. However, these manually intensive methods are low-throughput and the measurements for antibacterial activity can be subjective. Given these constraints, this study developed a platform to greatly increase the isolation and screening of coral probiotics. Using the new platform, microbial cells were isolated from a coral mucus sample using a microfluidic cell sorter, and isolates were then screened for inhibitory activity against target pathogenic bacteria modified to express yellow fluorescent protein (YFP) that allows growth quantification using a microplate reader. In a single run using the platform, 433 isolates were sorted then individually screened against two target pathogens within six days. The workflow described herein represents a proof of concept for high-throughput environmental probiotic discovery with potential to push forward treatment development for SCTLD and future disease outbreaks.

## INTRODUCTION

Coral reefs, among the most biodiverse ecosystems on the planet, are increasingly threatened by anthropogenic stressors that drive their global degradation (Beck et al., 2018; Ferrario et al., 2014; Graham and Nash, 2013; Ortiz and Tissot, 2012; Spalding et al., 2017; Teh et al., 2013). These stressors often exacerbate disease outbreaks, causing irreparable damage; in the Caribbean, a hotbed for coral disease (Weil, 2004), region-wide outbreaks have caused live coral cover to decline by as much as 80% in some areas (Gardner et al., 2003). A recent outbreak of stony coral tissue loss disease (SCTLD) has rapidly spread throughout all of Florida’s Coral Reef and at least 28 Caribbean countries and territories since its emergence in 2014 (Atlantic and Gulf Rapid Reef Assessment, n.d.; Estrada-Saldívar et al., 2020; Precht et al., 2016; Walton et al., 2018). SCTLD causes tissue to slough off of the coral, which can result in complete mortality within weeks of infection (Aeby et al., 2019; Florida Department of Environmental Protection, 2018). Impacting over 30 stony coral species (Atlantic and Gulf Rapid Reef Assessment, n.d.), SCTLD poses a dire threat to critical reef-building species, whose loss threatens the ability of these ecosystems to provide essential ecological and economic services (Estrada-Saldívar et al., 2020).

The causative agent(s) for SCTLD remain unknown, however, pathogenic bacteria play a role as disease progression slows or stops in response to antibiotics (Aeby et al., 2019). While a broad-spectrum antibiotic paste is the dominant field treatment for SCTLD, antibiotics cannot prevent reinfections (Shilling et al., 2021) and risk producing antibiotic-resistant pathogens. In response, Ushijima et al. (2023) found a probiotic, *Pseudoalteromonas* sp. strain McH1-7, originating from the microbiome of a susceptible coral species, *M. cavernosa*, that did not develop disease signs after exposure (Ushijima et al., 2023). Probiotics are defined by the World Health Organization as “live microorganisms which when administered in adequate amounts confer a health benefit on the host” (Joint FAO/WHO Expert Consultation, 2001). The bacterium, strain McH1-7, displayed broad-spectrum antibacterial activity and was found to stop or slow disease progression on 62% of corals in lab trials (Ushijima et al., 2023). Furthermore, field applications of strain McH1-7 also slowed or stopped disease progression on treated colonies (Pitts et al., 2025), further supporting the viability of harnessing the coral microbiome to treat SCTLD.

Although strain McH1-7 has demonstrated viability as an SCTLD intervention method *in situ*, a single probiotic strain may not be effective across coral species, many of which feature more genetic diversity than previously described (Grupstra et al., 2024). In addition, it is also critical that probiotic candidates are isolated from the native environment where they will be applied in order to offset the risk of introducing foreign strains with unknown ecological impact (Peixoto et al., 2022). Given these considerations, a diverse portfolio of probiotics should be sourced from multiple species across impacted regions to develop treatments across the wide host range and geographic extent of SCTLD. An expansive library of probiotic candidates could therefore provide a valuable tool for managing the spread of SCTLD and mitigating devastating impacts on coral reef ecosystems.

Developing a robust pipeline of probiotic candidates requires developing high-throughput methods that will rapidly scale up each step of the discovery process from sourcing isolates to screening them for activity. However, the development of probiotic-based treatments is currently hampered by labor-intensive isolation, purification, and screening methods. For example, isolating strain McH1-7 involved spreading dilutions of homogenized coral samples on agar plates, then manually picking and purifying colonies before inoculating into liquid media for assays (Ushijima et al., 2023). Similar methods are employed in probiotic discovery for addressing coral and other wildlife stressors, but these approaches are often low-throughput (Fragoso ados Santos et al., 2015; Hoyt et al., 2015; Iorizzo et al., 2022; Rosado et al., 2019; Santoro et al., 2021; Truong et al., 2023). The highest published number of microbes isolated and purified at one time for coral probiotic research is 133 strains (Santoro et al., 2021). Furthermore, screening for antibacterial activity using common methods presents scalability challenges. For example, the commonly used spotting assay consists of manually spotting test microorganisms on to an agar plate spread with pathogen/target culture, and measuring the clear zone (zone of growth inhibition) that forms after incubation, which is an indicator of the magnitude of inhibition towards the pathogen (Fragoso ados Santos et al., 2015; Rosado et al., 2019; Santoro et al., 2021; Zhang et al., 2021). Although this simple, low-cost assay helped identify strain McH1-7 (Ushijima et al., 2023), the process is time-consuming, and results can be subjective. Other bioassays for antibacterial activity, including disk diffusion, agar well diffusion, broth or agar dilution, and cross streak method, share similar limitations in terms of time, labor, and scalability (Balouiri et al., 2016).

To address these bottlenecks and accelerate the probiotic discovery process, we transformed conventional benchtop microbiology techniques into high-throughput methods. Starting with the initial step of isolating and purifying microbes from coral samples for downstream screening, we used microfluidic cell sorting to replace manual colony picking. This technology has already shown potential in rapidly isolating bacterial cells without compromising cell integrity, allowing for improved DNA recovery compared to other cell sorting technologies (Wiegand et al., 2021). Based on its efficiency, minimal manual input, and gentle cell handling, microfluidic cell sorting was an effective replacement for labor-intensive procedures commonly used to isolate and purify coral probiotics. In addition, the screening process for antibacterial activity was automated to address challenges related to scalability and measurement accuracy. To this end, we converted the screening assay into a liquid-based assay formatted to a microwell plate. For this assay, we modified target pathogens (referred to as the “target strains”) to constitutively express fluorescence, allowing changes in fluorescence to quantitatively indicate target strain growth when co-cultured with potential probiotic candidates (referred to as the “tester strains”). This format allowed the inhibition metric to be quantified by a microplate reader, overcoming issues with subjectivity in the commonly used screening assay. Without a known primary agent(s) for SCTLD, the target strains were potentially opportunistic strains isolated from coral tissue lesions (Ushijima et al., 2023) and are from bacterial families enriched at SCTLD lesions (Meyer et al., 2019; Rosales et al., 2020). Similar liquid-based assays have been successfully used in other contexts, such as testing antituberculosis agents against *Mycobacterium* strains (Collins et al., 1998) and probiotic strains against *Vibrio harveyi* (Pham et al., 2014). Overall, the discovery of promising probiotic candidates can be significantly accelerated by deploying automation and developing a recombinant microbial assay.

As SCTLD continues to spread, there is an urgent need to rapidly find and deploy probiotic candidates on surviving colonies. This paper presents a proof-of-concept for a high-throughput probiotic discovery platform capable of isolating, culturing, and screening probiotic candidates with greater efficiency and capacity than conventional methods. By incorporating a microfluidic single cell dispenser and developing a recombinant fluorescence-based screening assay, this platform aims to address critical bottlenecks in advancing probiotic discovery to combat SCTLD. Furthermore, the flexibility of the platform to isolate cells from any liquid substrate and include any plate-based assay allows for its future application beyond coral probiotic discovery.

## METHODS

### Bacterial strains and growth conditions

The bacterial strains used for this study are listed in Supplementary Table S1. Marine bacteria were grown in Tris-buffered glycerol artificial seawater broth (GASW-Tris) (Ushijima and Häse, 2018). GASW-Tris consisted of 20.8 g. NaCl, 0.56 g KCl, 4.8 g MgSO_4_·7H_2_O, 4.0 g MgCl_2_·6H_2_O, 0.01 g K_2_HPO_4_, 0.001 g FeSO_4_·7H_2_O, 2.0 g Instant Ocean Sea Salts, 6.03 g Tris Base (C_4_H_11_NO_3_), 4.0 g tryptone, 2.0 g yeast extract, and 2.0 mL glycerol into one L of ddH_2_O. The mixture was then adjusted to a pH of 8.0 using concentrated HCl prior to autoclaving. For solid media, 15 g/L of agar was added before autoclaving. The wash buffer, Tris-buffered artificial seawater (ASW-Tris), was made using the same recipe as GASW-Tris but without tryptone, yeast extract, and glycerol. All marine bacteria were grown at 28.5°C.

For conjugation, marine bacteria were grown on GASW-Tris agar. Antibiotics for marine bacteria were used at the following final concentrations: chloramphenicol (OfT6-21: 5 µg/µL; McT4-56: 10 µg/µL). All *E. coli* strains were grown in Lysogeny broth (LB) (Miller formulation with final concentration of 1% NaCl) with 100 µg/µL (final concentration) diaminopimelic acid (DAP) added to liquid media for auxotrophic strains. Antibiotics for *E. coli* used for conjugations were used at the following concentrations: kanamycin (50 µg/µL) and chloramphenicol (15 µg/µL). Chromogenic *E. coli* strains were cultured in LB (Miller) supplemented with chloramphenicol (15 µg/µL). All *E. coli* strains were grown at 37°C unless otherwise stated.

### Sample collection and processing

Corals were collected using SCUBA around Florida and housed at the Smithsonian Marine Station. Sample fragments from various apparently healthy colonies were provided by Smithsonian staff for this project. A list of the coral colony IDs and collection locations are provided in Supplementary Table S2. Coral fragments were then transported to University of North Carolina Wilmington (UNCW) in 250 mL of filtered seawater (FSW) in a cooler bag. The FSW was filtered down to 0.22 µm and cycled through a UV-sterilization system to eliminate microbial contamination as previously described (Ushijima et al., 2023). Upon arrival at UNCW, the coral fragments were immediately processed. The bottles containing the coral fragments from Florida were placed on an orbital shaker (Orbi-Shaker XL shake table, Benchmark) set to 90 rpm for ∼45 minutes at room temperature (approximately 20°C). The FSW was then filtered through a sterile 5 µm-pore syringe (Acrodisc 32 mm syringe filter with 5 µm Supor Membrane (polyethersulfone), 4650, PALL) to exclude large debris like coral skeleton fragments. Then, the FSW filtrate was filtered through a sterile 45 mm 0.22 µm-pore membrane (Nitrocellulose membrane, SA1J789H5, Merck Millipore) using an autoclave-sterilized vacuum filtration system. The membrane was then vortexed in 10 mL of autoclave-sterilized ASW-Tris in a sterile 50-mL conical for approximately 3 minutes to resuspend the captured microbes. Then, 1 mL aliquots were mixed with sterile glycerol in ASW (20% v/v glycerol final concentration) and cryopreserved at -80°C.

Additionally, mucus was collected from a *Montastraea cavernosa* coral colony with no gross disease signs (tissue loss lesions or bleaching) from Montserrat (mucus collection details listed in Supplementary Table S2). The collection was done by drawing up mucus using a 10 mL sterile needleless syringe while gently agitating the coral’s surface but not causing tissue damage. Multiple syringe samples were taken from the same coral to fill a sterile 15 mL conical with mucus. The mucus sample was transferred into a sterile conical tube. The conical was then placed in a Ziploc bag wrapped in paper towels, placed in a foam box with ice packs, and then shipped overnight to UNCW. Upon arrival at UNCW, the mucus sample was stored at 4°C until it was processed, which occurred within 8 h of arrival. To collect the microbes from the samples, ∼12 mL of the mucus was filtered through a sterile 0.22 µm membrane (Nitrocellulose membrane, SA1J789H5, Merck Millipore) using an autoclave-sterilized vacuum filtration system. Then, the membrane was transferred to a sterile 50-mL conical filled with ∼6.5 mL of autoclave-sterilized ASW using sterile forceps. The conical was vortexed for 3 minutes then 1 mL aliquots were mixed with sterile glycerol in ASW (20% v/v glycerol final concentration) and cryopreserved at -80°C.

### Plasmid construction and conjugation

All plasmids used in this study are listed in Supplementary Table S3 and all DNA oligonucleotides and their sequences are listed in Supplementary Table S4. To track the growth of marine bacteria, a replicative YFP expression vector was created. All plasmids were constructed using *E. coli* strain DH5α MCR (Grant et al., 1990). First, the replicative vector pBU115 was created by amplifying pEVS78 (Stabb and Ruby, 2002) with the primer set pEVS78-SmaI-F and pEVS78-SmaI-R. The resulting PCR product was digested with *Dpn*I and *Sma*I before being self-ligated. The resulting vector has a *Sma*I site introduced between the MCS and the natural *Hind*II/*Cla*I site.

The vector pBU159 is pBluescriptSK+ (Stratagene) with *yfpmut2* cloned in it using the *Eco*RI and *Pst*I sites. The primers yfpmut2-EcoRI-up-F and pKL183-yfp-PstI-R were used to amplify *yfpmut2* from the vector pKL183 (Lemon and Grossman, 2000) and introduce a unique *Eco*RV site upstream of *yfpmut2.* The product was then digested with *Eco*RI and *Pst*I and then cloned into pBluescriptSK+ previously linearized with the same enzymes. Blue/white screening and PCR using the primers yfpmut2-EcoRI-up-F and M13R were used to screen colonies. The YFP gene does not have a promoter in this vector. There is a unique *Eco*RV site upstream and a unique *Sma*I site downstream of *yfpmut2*. The vector pBU160 is pBU159 with the *bla* promoter (P*_bla_*) introduced upstream of *yfpmut2*. The *bla* promoter was amplified from pBBR1MCS-4 (Kovach et al., 1994) with the primers bla-SmaI-F and bla-SmaI-R. The 1113 bp PCR product was digested with *Sma*I and cloned into the *Eco*RV site of pBU159. Colonies were screened for fluorescence (503 nm excitation, 524 nm emission). The vector pBU164 is a constitutive YFP expression vector with *yfpmut2* driving by P*_bla_* cloned into pBU115. Using the primers M13-F and pKL183-yfp-PstI-R, P*_bla_*-*yfpmut2* was amplified from pBU160. The PCR product was cloned into the *Sma*I site of pBU115. Colonies were screened for fluorescence and using PCR with the primers pEVS78-MCS-F and pEVS78-MCS-R. The final vector, pBU164, was then transformed into *E. coli* strain β3914 (Le Roux et al., 2007) for conjugations.

The conjugation of marine bacteria (*Leisingera* sp. strain McT4-56 and *Vibrio coralliilyticus* strain OfT6-21) was performed as bi-parental conjugations with *E. coli* strain β3914 (Le Roux et al., 2007). One mL of 15 h GASW-Tris cultures of a marine strain were washed with ASW-Tris three times and resuspended in 25 µL of ASW-Tris. One mL of a β3914 culture with DAP was washed three times with PBS and resuspended in 25 µL of ASW-Tris. The *E. coli* and marine bacterium were gently mixed 1:10 (*Vibrio*:*E. coli*), and then 15 µL aliquots were spotted on GASW-Tris agar that previously had 30 µL of DAP stock (stock solution at 100 µg/mL) spread on it. The conjugations spots were then incubated for 24 h at 28.5°C. After incubation, spots were scraped off the plate and combined, washed three times with ASW-Tris, and then resuspended in one mL of ASW-Tris. For conjugations with *Leisingera* sp. strain McT4-56, dilutions were spread on GASW-Tris agar with 10 µg/µL of chloramphenicol. For conjugations with *V*. *coralliilyticus* OfT6-21, dilutions were spread on GASW-Tris agar with 5 µg/µL chloramphenicol. Plates were incubated at 28.5°C for 48 h. Colonies were streaked out for purification then inoculated into GASW-Tris broth with chloramphenicol (10 µg/µL for McT4-56; 5 µg/µL for OfT6-21). The cultures were checked for fluorescence (503 nm excitation, 524 nm emission) using a microplate reader (Varioskan Lux, Thermo Fisher Scientific), as well as with PCR using the primers pEVS78-MCS-F and pEVS78-MCS-R. Liquid cultures of the most fluorescent strains were mixed with sterile glycerol in ASW (20% v/v glycerol final concentration) and cryopreserved at -80°C.

### Determining the effect of dilution on microbial growth in coral mucus

The coral mucus from eight coral fragments across three species (coral details in Supplementary Table S2) was thawed from cryopreservation at room temperature (approximately 20°C) for 10 minutes. After thawing, each sample was serially diluted (2^-1^ to 2^-5^) with autoclave-sterilized ASW-Tris. For each coral mucus sample, 150 µL of each dilution was mixed with 50 µL of GASW-Tris broth in a sterile clear flat bottom 96-well microplate (Tissue Culture Plate, non-treated, sterile, 10861-562, VWR) in triplicate. Each microplate also included the undiluted coral mucus of each sample mixed with 50 µL of GASW-Tris broth in triplicate. In addition, the microplate included wells with 150 µL ASW and 50 µL GASW-Tris broth as media blanks in triplicate. The plate was then incubated in a plate reader (Varioskan Lux, Thermo Fisher Scientific) at 28.5°C with 120 rpm continuous shaking speed with force set to medium (name of setting on plate reader) while the optical density measured as absorbance at 600 nm (OD_600nm_) was measured every hour for 24 h.

### Determining the purity of single-cell cultures created by the single cell dispenser

To assess the accuracy of the single cell dispenser (BF.sight, CYTENA), five chromogenic *E. coli* strains (Supplementary Table S1) were revived from cryopreservation on LB (Miller) plates with chloramphenicol (15 µg/µL). One colony of each strain was then inoculated into 1 mL LB (Miller) broth with chloramphenicol and cultured at 37°C overnight. Each culture was then pelleted by centrifugation (8,000 rpm for 2 min, Microfuge 20, Beckman Coulter) and washed twice with autoclave-sterilized 1x PBS. Each pellet was then resuspended in 1 mL 1x PBS and diluted to an optical density measured at 600 nm (OD_600nm_) of 0.9 – 1.0. Then, 100 µL of each culture was combined then pelleted by centrifugation (8,000 rpm for 2 min, Microfuge 20, Beckman Coulter) and resuspended in 5 mL of sterile 1x PBS. To create pure *E. coli* cultures from this mixture, 50 µL diluted 1:10^4^ using sterile 1x PBS was added into a sterile microfluidic cassette (Cartridge, CY.XS.CAR.013, CYTENA) in the microfluidic single cell dispenser (BF.sight, CYTENA). Single cells were identified by the dispenser and sorted into a 96-well microwell plate (Cell culture microplate, PS, F-bottom, 655086, Greiner Bio-One) filled with LB (Miller) broth with chloramphenicol (15 µg/µL). After sorting, the plates were incubated at 37°C for 48 h then placed at room temperature (approximately 20°C) for four days, after which each well with visible growth was streaked out on to LB (Miller) agar plates supplemented with chloramphenicol (15 µg/µL) and incubated at 37°C for 48 to 64 h for pigmentation development.

### Determining the fluorescence of the modified target bacteria in co-culture

To determine how each target strain responds to co-culturing, the two target strains were revived from cryopreservation on GASW-Tris agar with chloramphenicol at their respective concentrations (OfT6-21: 5 µg/µL; McT4-56: 10 µg/µL). Ten biological replicates of strain OfT6-21 with pBU164 were each inoculated into 2 mL GASW-Tris broth with chloramphenicol and grown at 28.5°C on an orbital shaker set to 75 rpm for 12 h. Each culture was then pelleted by centrifugation (8,000 rpm for 2 min, Microfuge 20, Beckman Coulter) and washed with autoclave-sterilized ASW three times. Each pellet was then resuspended in 1 mL GASW-Tris and diluted to an OD_600nm_ of 0.8 – 0.9 using autoclave-sterilized ASW. In addition, ten biological replicates of the putative coral probiotic bacterium strain McH1-7 (Ushijima et al., 2023) and of each target strain’s wildtype (OfT6-21 and McT4-56) were grown in 2 mL of GASW-Tris broth at 28.5°C for 12 hours. Each culture was then diluted with autoclave-sterilized ASW to an OD_600nm_ of 0.8 – 0.9.

Then, the co-cultures were prepared by inoculating 10 µL of the target strain and 10 µL of strain McH1-7 (*n* = 10) into 180 µL of GASW-Tris broth per well in a black 96-well plate (Cell culture microplate, PS, F-bottom, 655086, Greiner Bio-One) (*n* = 10 per target strain). Additionally, 10 µL of each target strain and 10 µL of its respective wildtype strain was inoculated into 180 µL of GASW-Tris broth per well into the same black flat bottom 96-well plate (Cell culture microplate, PS, F-bottom, 655086, Greiner Bio-One) (*n* = 10 per target strain). Each co-culture was prepared in triplicate. To measure the RFU of the target strain in monoculture, 10 µL of each target strain was also inoculated into 180 µL of GASW-Tris broth in triplicate. To equalize the volumes of each well in the microwell plate, 10 µL of autoclave-sterilized ASW was added to the wells inoculated with cultures of the target strain so the total volume of all the wells was 200 µL. Wells with only GASW-Tris broth were included in triplicate as a media blank for background fluorescence. The plate was covered with a sterile sealing membrane (Breathe Easy Sealing Membrane, 70536-10, Electron Microscopy Sciences) and incubated at 28.5°C with 120 rpm shaking at medium force in the plate reader (Varioskan Lux, Thermo Fisher Scientific) with an RFU measurement (503 excitation, 524 emission) and OD_600nm_ taken every hour for up to 48 h for strain McT4-56 co-cultures and every 10 min for up to 24 h for strain OfT6-21 co-cultures.

### Isolating, culturing, and screening coral microbes using the high-throughput platform

To isolate and screen microbes for antibacterial activity in a high-throughput manner, microbes captured from the mucus of a healthy *Montastraea cavernosa* coral (coral details in Supplementary Table S2) were recovered from cryo-preservation at room temperature (approximately 20°C) for 15 minutes. Then, 65 µL of the sample was added into a sterile microfluidic cassette (B.sight Dispensing Cartridge, CY.XS.CAR.013, CYTENA) in a microfluidic single cell sorter (BF.sight, CYTENA). The cassette was positioned in front of a video microscope that provided a real-time video feed and could detect the size and roundness of the cells in the sample. The sorting parameters on the BF.sight were adjusted to: Region of Interest (ROI) = 325, Detection Threshold = 1.0, Cell Size = 0 - 5 µm, Cell Roundness = 0 - 0.9. Piezo actuation of the cassette created microdroplets (∼50 pL) from the sample. Before sorting cells, the position of the microdroplet was tested by dispensing 300 droplets into three wells of a clear flat bottom 384-well plate (pureGRADE, clear, F, 781680, BRANDplates) without any media in the wells to ensure the droplet was positioned to dispense into the well. Once it was confirmed the droplet was dispensing into the well, the 384-well plate was replaced with a sterile clear 384-well flat bottom plate (pureGRADE, clear, F, 781680, BRANDplates) filled with 90 µL of GASW-Tris broth per well. When cell sorting was activated, single cells detected within the ROI were encapsulated within a microdroplet and dispensed into the well below. All microdroplets containing multiple cells or no cells were disposed of by a vacuum shutter. Each microwell plate had two wells without cells to serve as the media blank for an absorbance reading.

To culture the sorted cells, the plates were incubated at 28.5°C while shaking at 120 rpm (Orbi-Shaker, Gene Mate) for 48 h. After incubation, an OD_600nm_ measurement was taken using the plate reader to identify wells with growth. A robotic liquid handler (Liquid Handling Station Flow, BRAND) was programmed to cherry pick from wells with measurable growth and inoculate 10 µL of each culture into wells with 150 µL of GASW-Tris broth in a sterile clear flat bottom 96-well plate (Tissue Culture Plate, non-treated, sterile, 10861-562, VWR). The inoculated plates were then incubated at 28.5°C while shaking at 120 rpm (Orbi-Shaker, Gene Mate) for 48 h.

To screen the isolate cultures for antibacterial activity against putative coral pathogens, the two target strains were revived on GASW-Tris agar with their respective antibiotic concentrations (OfT6-21 with pBU164: 5 µg/µL; McT4-56 with pBU164: 10 µg/µL). The coral probiotic strain McH1-7 and each target strain’s wildtype strain were revived on GASW-Tris agar. Then, each target strain was inoculated into 5 mL of GASW-Tris broth with their respective antibiotic concentrations. All other strains were inoculated into 5 mL of GASW-Tris broth. All the strains were cultured for up to 23 h at 28.5°C on a tube rotator set to 65 rpm. Then, 2 mL of each target strain culture was pelleted by centrifugation (8,000 rpm for 2 min, Microfuge 20, Beckman Coulter) and washed with autoclave-sterilized ASW three times then resuspended in 2 mL of GASW-Tris broth. Each culture was then adjusted to an OD_600nm_ of 0.8 –0.9 using autoclave-sterilized ASW.

To screen for antibacterial activity against the target strains, each tester culture (coral mucus isolate) was co-cultured with each target strain separately in a sterile black flat bottom 96-well plate (Cell culture microplate, PS, F-bottom, 655086, Greiner Bio-One). All liquid transfers for the antibacterial screen were conducted by a robotic liquid handler. First, 10 µL of each target strain culture was inoculated into 180 µL of GASW-Tris broth in the microwell plate. Then, 10 µL of each tester culture, including the positive control (strain McH1-7) was inoculated into each well containing a target strain. Each plate also included two wells of the target strain in monoculture as a no-treatment control and two wells with only GASW-Tris broth as a control for background fluorescence. The plates were incubated at 28.5°C with shaking at 120 rpm (Orbi-Shaker, Gene Mate). After 7 hours of culturing, the RFU (503 excitation, 524 emission) of each co-culture was measured using the plate reader (Varioskan Lux, Thermo Fisher Scientific).

The inhibitory activity of each tester strain was determined by the reduction in target strain RFU in the co-culture. Cryo-stocks were made from tester strains with strong (reduced target strain RFU by >80%) and weak (reduced target strain RFU by <50%) inhibitory activity using sterile glycerol diluted with ASW (20% v/v glycerol final concentration) and stored at - 80°C.

### Re-testing tester strains identified by the high-throughput platform screen

To confirm the activity of the coral mucus isolates identified as having strong or weak activity by the high-throughput assay, isolates with strong (*n* = 10 for both target strains) and weak (*n* = 4 for strain McT4-56 and *n* = 10 for strain OfT6-21) inhibitory activity were revived from cryopreservation and screened against the target strain it inhibited in the high-throughput screen previously performed (see above). First, each isolate was revived on GASW-Tris agar and re-plated three times for purification. The target strains OfT6-21 with pBU164 and McT4-56 with pBU164 were revived on GASW-Tris agar with their respective antibiotic concentrations (OfT6-21 with pBU164: 5 µg/µL; McT4-56 with pBU164: 10 µg/µL) then inoculated into 2 mL of GASW-Tris broth with their respective antibiotic concentrations. All the other isolates were inoculated into 1 mL of GASW-Tris broth. All the cultures were incubated for 24 h at 28.5°C on a tube rotator set to 65 rpm. After culturing, the target strains were pelleted by centrifugation (8,000 rpm for 2 min, Microfuge 20, Beckman Coulter) and washed three times with autoclave-sterilized ASW. Then the pellets were resuspended in 2 mL of GASW-Tris broth and diluted to an OD_600nm_ of 0.8 – 0.9 using autoclave-sterilized ASW.

For the screening assay, a sterile black 96-well flat bottom plate (Cell culture microplate, PS, F-bottom, 655086, Greiner Bio-One) was filled with 180 µL of GASW-Tris broth per well. Then, 10 µL of the target strain was added to each well followed by 10 µL of the tester strain in triplicate using a robotic liquid handler. Each tester strain was screened against the target strain it inhibited in the high-throughput screen. The plate was then incubated at 28.5°C in the plate reader (Varioskan Lux, Thermo Fisher Scientific) shaking at 120 rpm at medium force (name of setting on plate reader) while the RFU (503 excitation, 524 emission) was measured every 10 min for strain OfT6-21 with pBU164, and hourly for strain McT4-56 with pBU164.

### Statistical analysis

To compare microbial growth across dilutions of coral mucus, the OD_600nm_ for eight coral mucus samples serially diluted twofold was tracked over time. The growth curve data was grouped by dilution, and the area under the curve (AUC) for all samples in each dilution was compared using a repeated measures one-way ANOVA to examine the effect of dilution on microbial growth (Field, 2018). Assumptions of normality were evaluated using a Q-Q plot and a Shapiro-Wilk test, and when found to be violated (Field, 2018), a Friedman test, followed by Dunn’s multiple comparisons test were used to analyze the effect of dilution on microbial growth (Fig. 1) (Dunn, 1964; Field, 2018).

**Figure 1.**
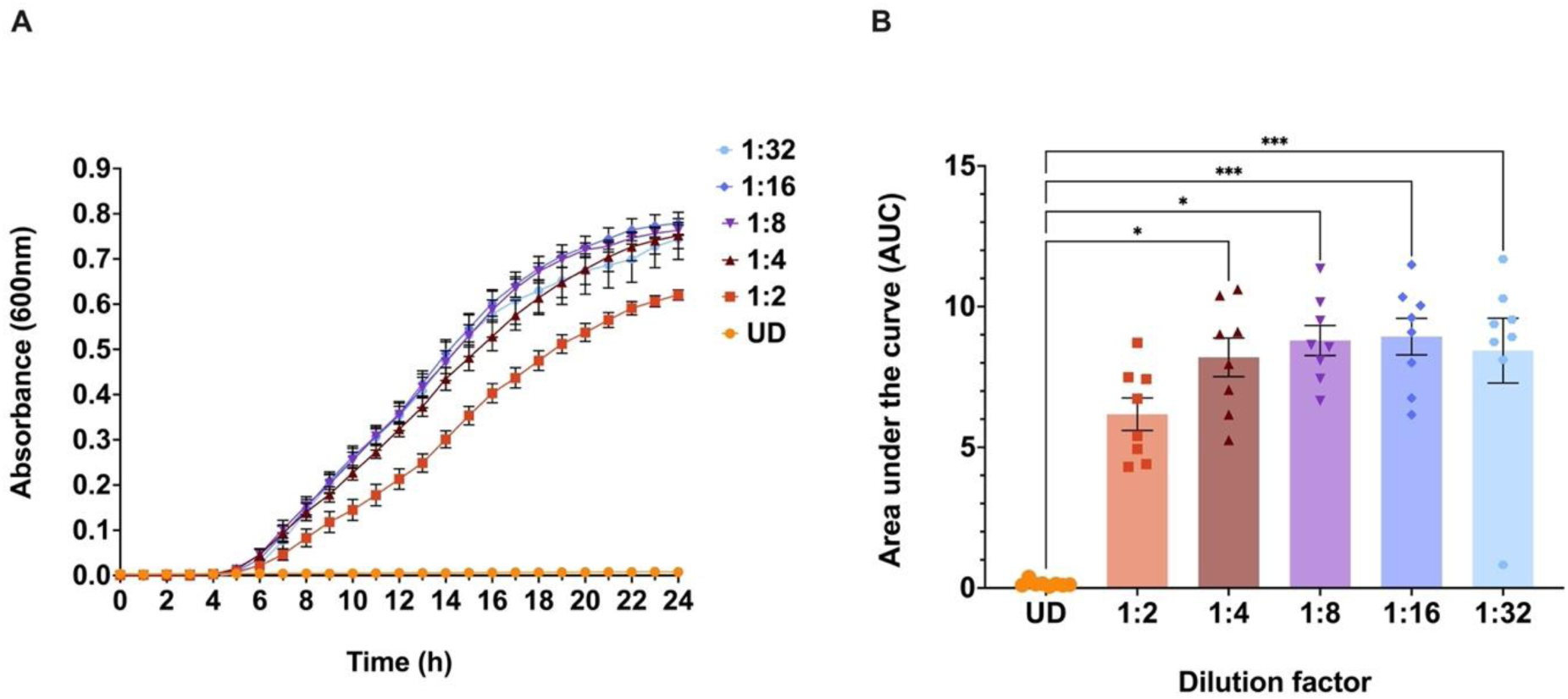
Total microbial growth in diluted and undiluted (UD) coral mucus samples. (A) Optical density at 600 nm (OD_600nm_) of coral mucus-associated microbes across a two-fold serial dilution series. The UD sample is represented by orange circles with subsequent dilutions shown as follows: 1:2 dilution (dark orange squares), 1:4 dilution (red triangles), 1:8 dilution (purple inverted triangles), 1:16 dilution (blue diamonds), and 1:32 dilution (blue hexagons). (B) Area under the curve (AUC) calculations for microbial growth across dilutions show significant differences compared to growth in the UD sample (χ^2^(5) = 28.29, *p < 0.0001, n* = 8). Asterisks indicate statistical significance (* = *p* ≤ *0.05, *** = p ≤ 0.001*). Error bars represent the standard error of the mean (SEM), and bar height in the bar graph represents the mean.

The growth of the target strains was measured by tracking RFU in monoculture (*n* = 10 for each target strain) and in co-culture (*n* = 10 each) over time, from which an AUC was calculated for each condition. An ordinary one-way ANOVA was used to compare target strain growth across three treatments (target strain monoculture, target strain + WT strain co-culture, target strain + McH1-7 co-culture) (Fig. 2C & 2D) (Field, 2018). These data were determined to be normally distributed as evaluated using a Q-Q plot and a Shapiro-Wilk test (Field, 2018).

**Figure 2.**
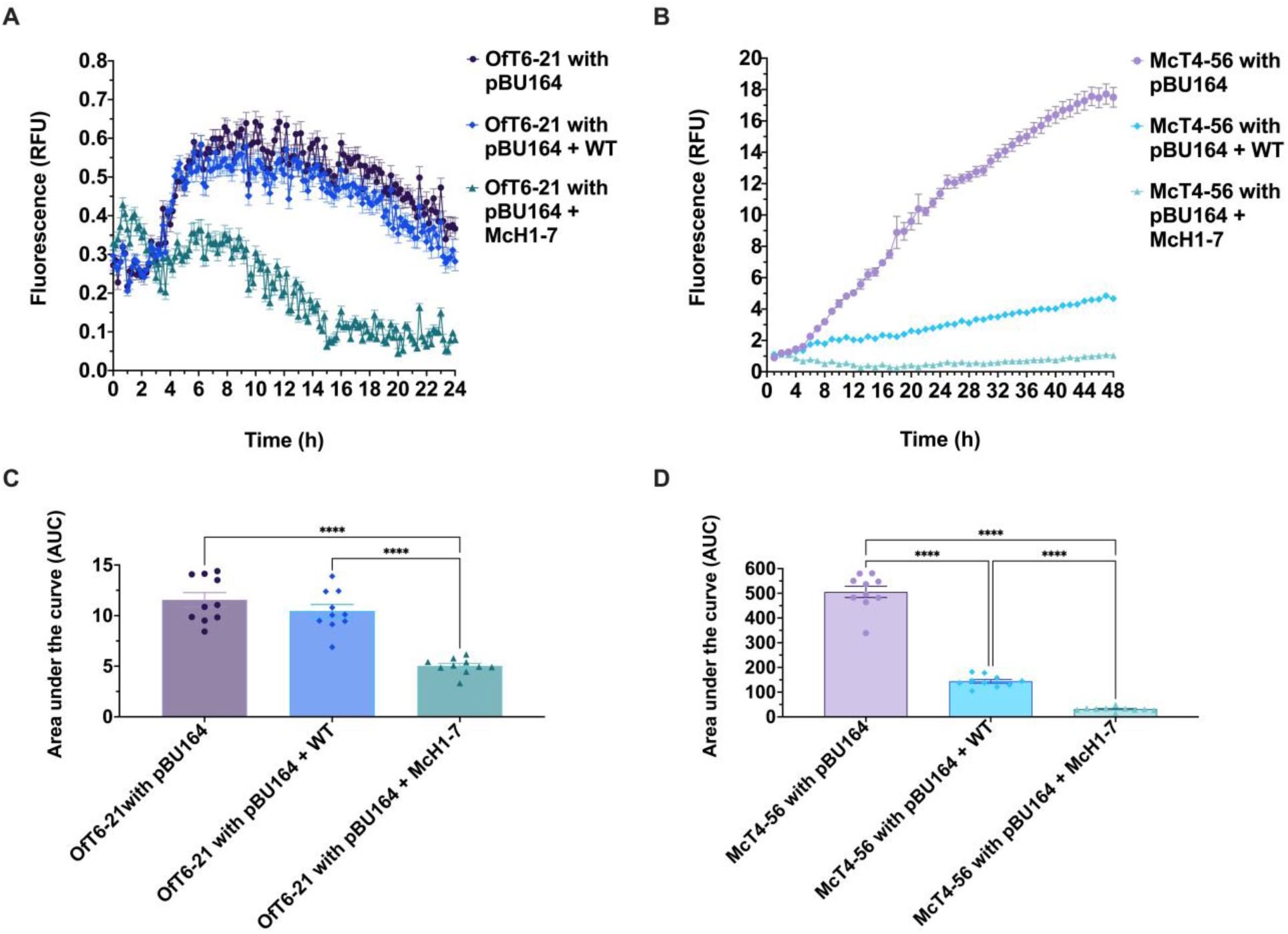
Comparison of target strain growth in monoculture and co-culture conditions. (A, B) Relative fluorescence units (RFU) over time for target strains OfT6-21 with pBU164 (A) and McT4-56 with pBU164 (B) cultured in GASW-Tris broth in a 96-well microtiter plate. Each target strain was grown in monoculture (purple line with circles), in co-culture with its respective wildtype (WT) strain (blue line with diamonds), and in co-culture with the putative coral probiotic, *Pseudoalteromonas* sp. strain McH1-7 (green line with triangles) (*n* = 10 per treatment). (B) Area under the curve (AUC) calculations for target strain OfT6-21 with pBU164 show significant differences in growth between the monoculture (purple circles), co-culture with its WT strain (blue diamonds), and co-culture with strain McH1-7 (green triangles) (one-way ANOVA; F(2, 27) = 37.90, *p < 0.0001*, *n* = 10 per group). (D) AUC calculations for target strain McT4-56 with pBU164 show significant differences in growth across monoculture (purple circles), co-culture with its WT strain (blue diamonds), and co-culture with strain McH1-7 (green triangles) (one-way ANOVA; F(2, 27) = 316.2, *p < 0.0001*, *n* = 10 per group). Asterisks indicate statistical significance (****= *p* ≤ *0.0001*). Error bars represent the standard error of the mean (SEM), and bar height in the bar graphs represent the mean.

To calculate the inhibitory activity of the tester strains screened through the high-throughput platform, the RFU of the target strain in each co-culture with a tester strain was compared to the average RFU of the target strain in monoculture on each microwell plate (Fig. 4). Tester strains that reduced the target strain RFU by >80% and <50% were identified as isolates with strong and weak inhibitory activity, respectively.

**Figure 3.**
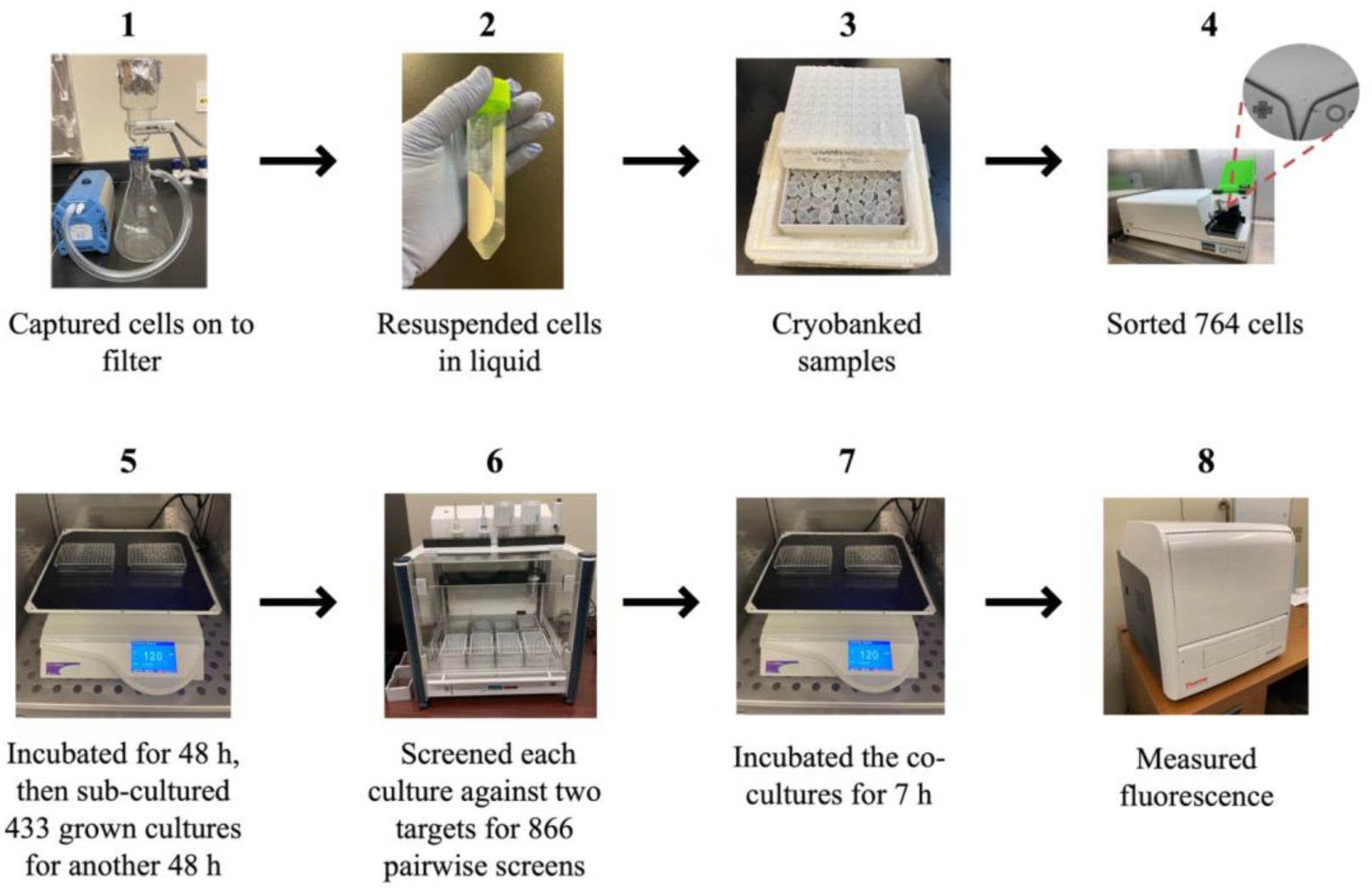
Workflow of the high-throughput platform with results from a single run. The platform was used to sort, culture, and screen microbes from a mucus sample collected from a healthy *Montastraea cavernosa* coral colony. Processing a sample through the entire workflow was completed within six days by one operator.

**Figure 4.**
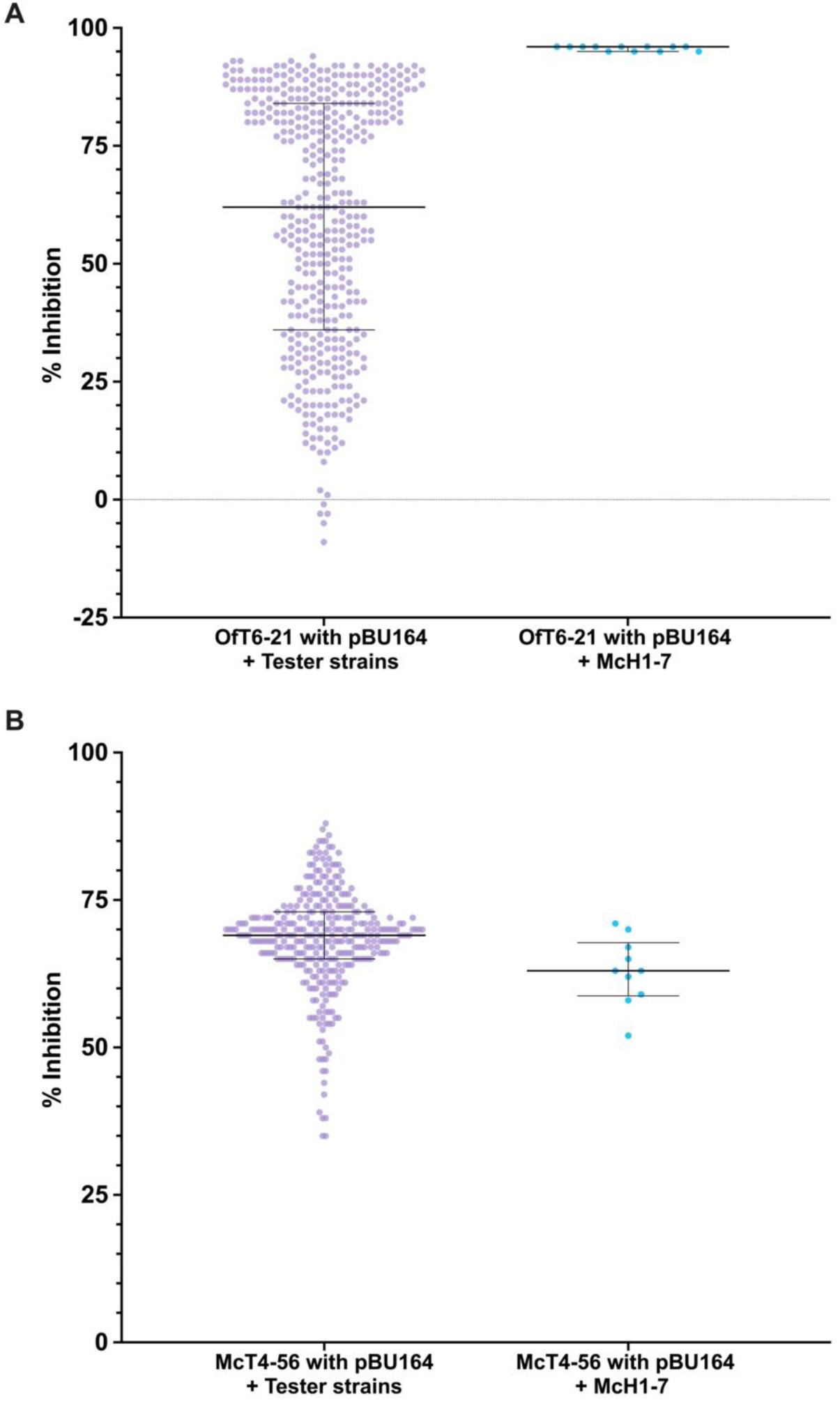
Inhibitory activity of coral mucus isolates (the tester strains) against target strains as measured in the high-throughput screen. (A, B) The inhibitory activity of each tester strain (*n* = 433 for OfT6-21 with pBU164; *n* = 345 for strain McT4-56 with pBU164; purple circles) was quantified as the percentage of target strain growth inhibition, calculated from RFU measurements. Target strains included strain OfT6-21 with pBU164 (A) and strain McT4-56 with pBU164 (B). For comparison, the screen also tested each target strain co-cultured with the known probiotic strain McH1-7 (blue circles). Lines represent the median, and bars represent interquartile ranges.

A subset of tester strains with either strong or weak inhibitory activity, as identified by the high-throughput screen, were re-tested against the target strains by tracking the RFU of the target strain in co-culture with each tester strain. The AUC for each growth curve was calculated and grouped as either inhibitory or non-inhibitory as determined by if they displayed strong or weak activity in the first high-throughput screen. An unpaired t-test was then used to compare inhibitory activity between the groups upon re-testing (Fig. 5) (Field, 2018). The data met assumptions of normality as evaluated using a Q-Q plot and a Shapiro-Wilk test (Field, 2018). All AUC calculations and statistical analyses were completed with GraphPad Prism version 10.0.3 (GraphPad Software, San Diego, CA, USA) (GraphPad Prism files in Supplementary File S1).

**Figure 5.**
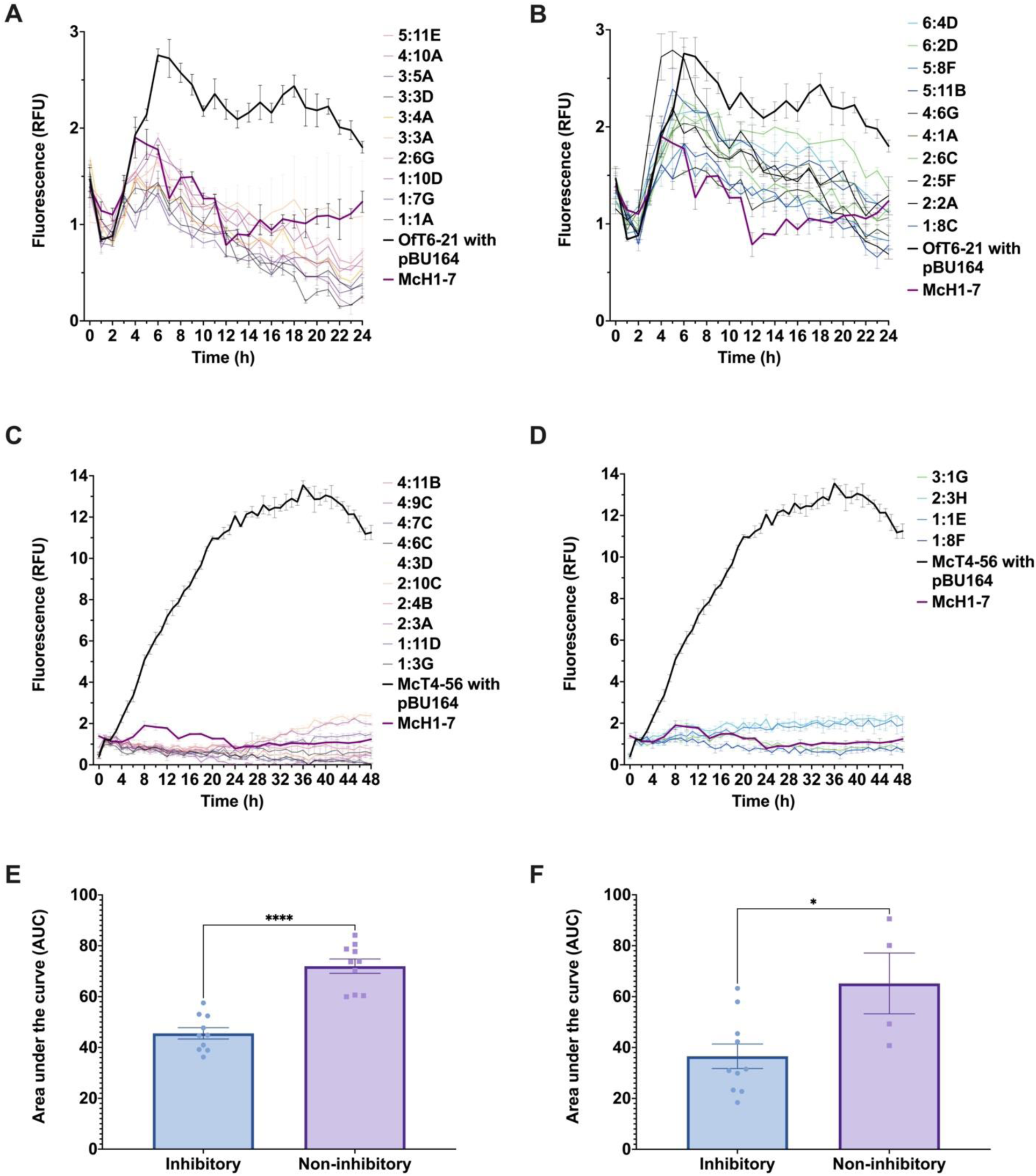
Re-testing inhibitory activity of tester strains isolated from coral mucus against target strains OfT6-21 with pBU164 and McT4-56 with pBU164. The following labels were used as identifiers for the tester strains: 5:11E, 4:10A, 3:5A, 3:3D, 3:4A, 3:3A, 2:6G, 1:10D, 1:7G, 1:1A, 6:4D, 6:2D, 5:8F, 5:11B, 4:6G, 4:1A, 2:6C, 2:5F, 2:2A, 1:8C, 4:11B, 4:9C, 4:7C, 4:6C, 4:3D, 2:10C, 2:4B, 2:3A, 1:11D, 1:3G, 3:1G, 2:3H, 1:1E, 1:8F. (A, B) The relative fluorescence units (RFU) of target strain OfT6-21 with pBU164 in co-culture with inhibitory isolates (A) (*n* = 10) and non-inhibitory isolates (B) (*n* = 10) tracked over time. (C, D) The RFU of target strain McT4-56 with pBU164 in co-culture with inhibitory isolates (C) (*n* = 10) and non-inhibitory isolates (D) (*n* = 4) tracked over time. The known probiotic strain McH1-7 (bolded purple line) and monoculture controls (bolded black line) are included for comparison (A – D). (E) Area under the curve (AUC) calculations for target strain OfT6-21 with pBU164 demonstrate significantly reduced growth in co-culture with inhibitory isolates compared to non-inhibitory isolates (unpaired t-test; *p < 0.0001*). (F) AUC calculations for target strain McT4-56 with pBU164 show a significant reduction in growth in co-culture with inhibitory isolates compared to non-inhibitory isolates (unpaired t-test; *p = 0.0186*). Asterisks indicate statistical significance (* = *p ≤ 0.05, **** = p ≤ 0.0001*). Error bars represent standard error of the mean (SEM), and bar height in the bar graphs represents the mean.

## RESULTS

### Diluting coral mucus microbes increases overall growth

To evaluate whether competitive interactions between coral mucus-associated microbes suppress total microbial growth in a sample, microbial cells collected from the mucus of eight coral fragments representing three species (*Orbicella faveolata*, *Siderastrea siderea*, *Montastraea cavernosa*) were serially diluted. The OD_600nm_ for each dilution, including the undiluted (UD) sample, was monitored over 24 h at 28.5°C (Fig. 1A). The AUC for each dilution of each sample was calculated, then grouped by dilution. As the normality assumption was violated (see Statistical Analysis section), a Friedman test was used to compare the AUC across dilutions. The analysis revealed a significant difference in growth across dilutions (χ^2^(5) = 28.29, *p < 0.0001, n* = 8) (Fig. 1B). Post-hoc analysis using Dunn’s multiple comparisons test demonstrated that microbial growth in the undiluted samples was significantly lower compared to all samples diluted by 1:4, 1:8, 1:16, and 1:32 dilution factors (UD vs. 1:4: *p = 0.0125*; UD vs. 1:8: *p = 0.0125*; UD vs. 1:16: *p = 0.0009*; UD vs. 1:32: *p = 0.0005*). These findings indicate that diluting coral mucus samples may alleviate competitive pressure among microbes, thereby increasing total microbial growth, independent of nutrient enrichment in the growth media. All raw data is available in Supplementary File S1.

### The single-cell dispenser accurately dispenses single cells to create pure cultures

The single-cell dispenser (BF.sight, CYTENA) was evaluated for its ability to isolate single cells into individual wells of a microtiter plate to create pure cultures. To test the accuracy of the dispenser, a mixed culture of chromogenic *E. coli* strains was sorted into 96-well plates. The resulting cultures were streaked onto agar plates to visually assess contamination by detecting if there were multiple chromogenic *E. coli* strains visible in each streak. Out of the 212 wells that exhibited growth, only two wells showed evidence of contamination (Supplementary Figure S2). These results demonstrate that the single-cell dispenser effectively isolates single cells into wells with a high level of accuracy, enabling the generation of pure cultures.

### Target strain RFU changes in co-culture based on tester strain inhibitory activity

The change in RFU emitted by a target strain when co-cultured with another isolate (tester strain) serves as a proxy for changes in target strain growth. To determine the impact of inhibitory activity, each target strain was co-cultured with its respective wildtype (WT) strain and separately with the putative coral probiotic *Pseudoalteromonas sp.* strain McH1-7 (Fig. 2). For target strain OfT6-21 with pBU164, analysis of the AUC for the RFU over time showed a significant difference across treatments (one-way ANOVA; F(2, 27) = 37.90, *p < 0.0001*, *n* = 10 per group) (Fig. 2C). Tukey’s post-hoc test showed that the AUC of the WT co-culture was significantly higher compared to the McH1-7 co-culture (*p < 0.0001*, 95% C.I. = [3.436, 7.419]) but not significantly different from the OfT6-21 with pBU164 monoculture (*p = 0.3666*, 95% C.I. = [-3.098, 0.8854]). Conversely, the McH1-7 co-culture exhibited a significantly lower mean AUC compared to the monoculture (*p < 0.0001*, 95% C.I. = [-8.526, -4.542]) (Fig. 2C).

For target strain McT4-56 with pBU164, RFU was also significantly different across treatments (one-way ANOVA: F(2, 27) = 316.2, *p < 0.0001*, *n* = 10 per treatment group) (Fig. 2D). Tukey’s post-hoc test indicated that the AUC for the McT4-56 with pBU164 monoculture was significantly higher than both its WT co-culture (*p < 0.0001*, 95% C.I. = [313.1, 410.9]) and its McH1-7 co-culture (*p < 0.0001*, 95% C.I. = [425.5, 523.3]). The WT strain’s inhibitory activity is likely attributable to resource competition, as the WT strain grows significantly faster than the genetically modified target strain (Supplementary Figure S1: Mann-Whitney U = 0, *p < .0022, n* = 6 per strain). However, strain McH1-7 was even more inhibitory, with the WT co-culture showing a significantly higher AUC than the McH1-7 co-culture (*p < 0.0001*, 95% C.I. = [63.52, 161.3]) (Fig. 2D).

These results suggest that strain McH1-7 exerts consistently stronger inhibition on both target strains compared to their respective WT strains. The strong inhibitory activity of strain McH1-7 is hypothesized to result from the production of broad-spectrum antibacterial compounds (Ushijima et al., 2023), whereas WT inhibition likely stems from resource competition. In summary, target strain RFU decreases in response to growth inhibition in co-culture with tester strains. All raw data is available in Supplementary File S1.

### Processing coral mucus isolates through the high-throughput platform

Microbes from the mucus of a healthy *M. cavernosa* coral colony were isolated, cultured, and screened for inhibitory activity against two target strains using the high-throughput platform. The entire process was completed within six days by one operator (Fig. 3). Following mucus processing (Fig. 3, steps 1 – 3), 764 cells were isolated, resulting in 433 individual cultures that were screened for antibacterial activity against the target strains (Fig. 3, steps 4 – 8).

The antibacterial activity assay included the following controls on each plate: target strain co-cultured with *Pseudoalteromonas* sp. strain McH1-7 (positive control), target strain monoculture (no-treatment control), and media blanks. While screening the isolates for antibacterial activity towards target strain McT4-56 with pBU164, a microtiter plate was placed incorrectly in the robotic liquid handler platform, leading to controls being added to the wrong columns. As a result, the 88 tester strains from that plate were excluded from analyses.

The inhibitory activity of each tester strain was determined by calculating the difference in endpoint RFU between the target strain monoculture and the target strain cultured with the tester strain (Fig. 4). The screening results revealed a much wider range of inhibitory activity against target strain OfT6-21 with pBU164 compared to strain McT4-56 with pBU164.

Interestingly, some tester strains appeared to enhance the growth of the target strain (Fig. 4A). All raw data is available in Supplementary File S1.

### Re-testing coral mucus isolates screened by the high-throughput platform

To evaluate the reproducibility of the high-throughput screening assay, a subset of tester strains spanning a range of inhibitory activity were revived and re-screened against the target strains. Before re-testing, the OD_600nm_ of the isolates and control strains were standardized to an OD_600nm_ of 0.8. The inhibitory activity of each tester strain was evaluated by tracking the fluorescence of the target strains over time in co-culture within a 96-well plate (Fig. 5A–D).

For target strain OfT6-21 with pBU164, the isolates previously identified as strongly inhibitory (*n* = 10) remained significantly more inhibitory compared to those identified as weakly inhibitory (*n* = 10) in the high-throughput screen (unpaired t-test: *p < 0.0001*) (Fig. 5E). Similarly, for target strain McT4-56 with pBU164, the strongly inhibitory isolates (*n* = 10) were still significantly more inhibitory than the weakly inhibitory isolates (*n* = 4) (unpaired t-test: *p < 0.0186*) (Fig. 5F). These findings demonstrate that the high-throughput screen still differentiates between isolates originally found to have strong or weak inhibitory activity upon re-testing. All raw data is available in Supplementary File S1.

## DISCUSSION

This study presents a proof-of-concept for a high-throughput workflow that addresses methodological roadblocks to create a scalable and efficient approach to isolate and screen probiotic candidates from environmental samples. In the first step of the workflow, labor intensive and low-yield conventional methods for probiotic isolation and purification were addressed by incorporating microfluidic cell sorting. In the run of the platform described herein, single cells were deposited into two 384-well microtiter plates, resulting in 433 (56.8%) wells with growth after 48 hours of incubation (Fig. 3). In contrast to manually purifying colonies, operator involvement was reduced to prepping well-plates with media, adjusting parameters on the cell sorter, and swapping plates after sorting was completed. To improve screening throughput, YFP-expressing target strains were incorporated into a liquid-based assay, enabling a plate reader to monitor target strain growth in competition assays using relative fluorescence units (RFU). In the run of the platform described in this study, 433 tester strains were screened against two target strains within a day using the fluorescence-based assay (Fig. 3). Besides improving throughput, performing competition assays using a plate reader eliminated the subjectivity associated with agar-based assays, which rely on manual measurements of inhibition zones with unclear margins and partial inhibition. The integration of high-throughput cell sorting and a liquid-based inhibition assay enabled a single operator to isolate and screen 433 tester strains against two target pathogens within five days (not including the time it took to process the coral mucus samples) (Fig. 3). Thus, the high-throughput platform demonstrates the potential to significantly accelerate and improve the precision of probiotic discovery from environmental samples.

The single-cell culturing conditions used in the platform may also provide improved growth conditions. Previous research has demonstrated that coral-associated microbes exhibit mutual antagonism, likely as a means to prevent the dominance of any single taxon (Ritchie, 2006; Rypien et al., 2010). However, coral probiotic studies typically rely on spreading coral tissue homogenate on to solid media to culture resident microbes (Rosado et al., 2019; Santoro et al., 2021; Ushijima et al., 2023). This approach may inadvertently limit the growth of certain taxa due to microbial competition. To investigate the impact of reduced competition on microbial growth, the growth of coral mucus microbes was tracked across varying dilutions, showing significantly more growth at higher dilutions (Fig. 1). Other studies have similarly addressed competition by encapsulating single bacterial cells in microdroplets to cultivate recalcitrant strains (Watterson et al., 2020; Zengler et al., 2005). This method is mimicked in the high-throughput platform, as single cells are also encapsulated in microdroplets that are then sorted into a microwell plate. The alleviation of competitive pressure afforded by single-cell culturing conditions can therefore potentially improve microbial cultivability.

In addition to significantly enhancing throughput, the cell isolation method used for this platform offers several advantages over commonly used methods. For example, single-cell isolation using dilution-to-extinction relies on correct cell concentration estimates and sample dilutions. Instead of relying on statistical inference or accurate dilutions, the cell sorter has an embedded camera that tracks cell movement to ensure that individual cells are isolated into each well of a microtiter plate. Microfluidic cell sorting also offers benefits over fluorescence-activated cell sorting (FACS). Though FACS has successfully isolated single microbial cells from marine environmental samples (Rinke et al., 2014), it requires labeling and applies shear forces, which may affect cell division. By avoiding these requirements, the microfluidic sorting method is particularly well-suited for isolating microbial cells for the development of clonal cultures.

Using a fluorescence-based method for a competition assay also offers significant benefits over other common methods, such as using optical density. To assess the inhibitory activity of a target strain using optical density, the optical density of the culture is measured after culturing with the supernatant of a tester strain (Chomwong et al., 2018; Kueneman et al., 2016; Woodhams et al., 2014). This method cannot assess cellular interactions as it does not distinguish between the growth of different strains in a mixed culture. In contrast, the RFU-based method used in the platform leverages the constitutive expression of YFP by the target strain to continuously measure target strain growth within a mixed culture (Fig. 2A and 2B). The fluorescence-based measurement accounts for cell-to-cell interactions over time, capturing the effects of competition across different growth phases (Fig. 2A and 2B). These advantages make the fluorescence-based assay a powerful tool for characterizing the inhibitory activity of potential probiotic candidates.

The fluorescence-based assay also provides a means for characterizing interactions beyond inhibitory activity. Specifically, fluorescence measurements could differentiate between modes of growth inhibition, such as resource competition versus direct antagonism (e.g., antimicrobial compound production) (Fig. 2). For example, there was a minimal change in RFU of target strain OfT6-21 with pBU164 when co-cultured with its wildtype (WT), suggesting resource competition. In contrast, when OfT6-21 with pBU164 was co-cultured with strain McH1-7, there was a significant decrease in RFU (Fig. 2A and 2C), indicating stronger inhibitory activity likely a result of more antagonistic interactions. As strain McH1-7 exhibits broad-spectrum antimicrobial activity, its strong inhibitory activity is likely due to the production of antimicrobial compounds (Ushijima et al., 2023). For the comparatively slower-growing target strain McT4-56 with pBU164, growth was also significantly inhibited by strain McH1-7 compared to its WT strain (Fig. 2B and 2D). However, unlike strain OfT6-21 with pBU164, strain McT4-56 with pBU164 exhibited a significant reduction in growth when co-cultured with its WT strain (Fig. 2B and 2D), likely due to differences in growth rate. The WT strain grew almost twice as fast as the engineered strain, highlighting the need to account for growth rate changes caused by plasmid introduction when developing target strains for fluorescence-based assays (Supplementary Figure S1). Overall, these findings highlight the ability of the fluorescence-based assay to shed light on the competitive dynamics between strains.

Broader trends in host-microbe interactions may also be elucidated through the fluorescence-based screening assay. For example, the strong inhibitory activity exhibited by the tester strains against the target strains (Fig. 4) may be attributed to their origin from a healthy coral colony devoid of SCTLD signs, suggesting enhanced defenses against SCTLD-associated pathogens. A future study comparing the inhibitory activity of microbes cultivated from healthy versus unhealthy corals could reveal differences in microbial defense mechanisms related to coral health state. Furthermore, the assay sheds light on pathogen colonization strategies within the coral microbiome. For example, the stronger inhibition of the tester strains towards strain McT4-56 (Fig. 4B), suggests that strain McT4-56 may depend on microbial dysbiosis caused by a prior disturbance to effectively colonize a microbiome. Overall, the fluorescence-based assay may enable deeper insights into both inhibitory mechanisms and pathogen dynamics compared to a traditional agar plate-based screen.

The high-throughput platform presents many opportunities for optimization. Beginning with the initial cell sorting step, using a live-cell-specific dye to differentially sort viable cells using fluorescent sorting could reduce the chances of sorting dead cells. Additionally, enriching the sample with growth media before sorting may maximize the number of live cells; however, this approach risks selecting for specific taxa. To mitigate this risk, an enrichment regime that preserves community diversity of a sample could be developed using 16S rRNA gene sequencing to evaluate the effects of media composition and enrichment duration on community structure. The diversity of sorted cells could also be improved by minimizing repeat sampling during sorting. While conventional methods for wildlife probiotic selection ensure microbial diversity among candidates by picking colonies with unique morphologies (Rosado et al., 2019; Santoro et al., 2021; Ushijima et al., 2023), such an approach cannot be done with cell sorting. To diversify the cells isolated by a sorter, different taxa can be selectively cultivated through tailored media formulations (Bomar et al., 2011). For example, as shown with human gut microbiota (Tamargo et al., 2018), adjusting media viscosity to better mimic coral mucus substrate may encourage the cultivation of diverse taxa.

The fluorescence-based screening assay can be further refined by multiplexing, allowing for co-culturing multiple target strains with each tester strain. This approach could assess broad-spectrum antibacterial capabilities and more accurately reflect biological interactions *in situ*. Individual strains within a mixed culture could be tracked by engineering target strains to express distinct fluorescent proteins with minimal spectral overlap (Schlechter et al., 2021).

A major benefit of the high-throughput platform is its adaptability: the workflow can be customized to sort from different liquid substrates and include any liquid-based biochemical assay. This methodological flexibility expands its application beyond coral probiotics to microbial characterization in other systems. Within the context of coral probiotic development, this platform enables the rapid identification of probiotics that enhance host fitness under various stressors in addition to disease. For instance, probiotic candidates that confer resistance to heat stress could be rapidly identified by incorporating thermal tolerance assays into the workflow. The modular design of the high-throughput platform thus supports the characterization of a wide range of probiotic candidates, offering significant potential to advance efforts aimed at mitigating threats to coral health.

## CONCLUSION

This study demonstrates the proof-of-concept for a high-throughput platform designed to accelerate the discovery of probiotic candidates for SCTLD treatment. Compared to conventional methods of probiotic isolation and screening, this platform reduced processing time while increasing throughput at every step: in a single run, over 700 isolates were obtained, and more than 400 pure cultures were screened for antibacterial activity against two target pathogens in parallel. Given that SCTLD has persisted in parts of the Caribbean for over a decade, the capacity of affected reefs to provide critical ecological services is at serious risk (Estrada-Saldívar et al., 2021, 2020; Toth et al., 2023). The survival and recovery of these valuable ecosystems will depend on novel solutions to mitigate damage to impacted reefs while safeguarding naïve ones. To this end, coral probiotics offer significant potential as a scalable, safe, and prophylactic approach to combat SCTLD. The methods outlined in this study represent a key step toward the broader adoption of microbiome-based interventions for SCTLD and future coral disease outbreaks.

## ACKNOWLEDGEMENTS

This work was supported by National Philanthropic Trust/Oceankind and the Florida Department of Environmental Protection (Award# B7F150) with funds awarded to B.U. as well as startup funds from UNCW to B.U. We thank Revive & Restore for project management with their Advanced Coral Toolkit program. We would also like to thank those involved in collecting coral mucus in Montserrat for this project including Dr. Greta Aeby, Emmy Aston and Andrew Myers of Island Solutions, and the Joint Nature Conservation Committee (JNCC). We also thank Dr. Valerie Paul and staff at the Smithsonian Marine Station at Fort Pierce, Florida for providing us with coral mucus samples.

## Data Availability Statement

The data generated or analyzed here are included in the article or are available from the corresponding author upon reasonable request.

## Funding

This work was supported by National Philanthropic Trust/Oceankind and the Florida Department of Environmental Protection (Award# B7F150) with funds awarded to B.U. as well as startup funds from UNCW to B.U.

## Conflicts of Interest

The authors declare no conflicts of interest.

## Notes

### Competing Interest Statement

The authors have declared no competing interest.

